# Analysis of Hippocampal Subfields in Sickle Cell Disease Using Ultrahigh Field MRI

**DOI:** 10.1101/2020.11.10.377564

**Authors:** Tales Santini, Minseok Koo, Nadim Farhat, Vinicius P. Campos, Salem Alkhateeb, Marcelo A. C. Vieira, Meryl A. Butters, Caterina Rosano, Howard J. Aizenstein, Joseph Mettenburg, Enrico M. Novelli, Tamer S. Ibrahim

**Affiliations:** Department of Bioengineering, University of Pittsburgh, Pittsburgh, PA, United States; Department of Electrical and Computer Engineering, University of São Paulo, São Carlos, SP, Brazil; Department of Psychiatry, University of Pittsburgh, Pittsburgh, PA, United States; Department of Epidemiology, University of Pittsburgh, Pittsburgh, PA, United States; Department of Radiology, University of Pittsburgh, Pittsburgh, PA, United States; Department of Medicine, University of Pittsburgh, Pittsburgh, PA, United States; Department of Pharmacology and Chemical Biology, University of Pittsburgh, Pittsburgh, PA, United States

## Abstract

Sickle cell disease (SCD) is an inherited hemoglobinopathy that causes organ dysfunction, including cerebral vasculopathy and neurological complications. Hippocampal segmentation with newer and advanced 7 Tesla (7T) MRI protocols has revealed atrophy in specific subregions in other neurodegenerative and neuroinflammatory diseases, however, there is limited evidence of hippocampal involvement in SCD. Thus, we explored whether SCD may be also associated with abnormalities in hippocampal subregions. We conducted 7T MRI imaging in individuals with SCD, including the HbSS, HbSC and HbS/beta thalassemia genotypes (n=37), and healthy race and age-matched controls (n=40), using a customized head coil. Both T1 and T2 weighted images were used for automatic segmentation of the hippocampus subfields. Individuals with SCD had significantly smaller volume of the Dentate Gyrus and Cornu Ammonis (CA) 2 and 3 as compared to the control group. Other hippocampal subregions also showed a trend towards smaller volumes in the SCD group. These findings support previous reports of reduced volume in the temporal lobe in SCD patients. Further studies are necessary to investigate the mechanisms that lead to structural changes in the hippocampus subfields and their relationship with cognitive performance in SCD patients.

## Introduction

Sickle Cell Disease (SCD) consists of a group of heterogeneous syndromes that share the inheritance of a mutated sickle hemoglobin (hemoglobin S, or HbS) [1]. SCD is one of the most common genetic disorders, with more than 5 million newborns carrying the mutated gene and more than 300,000 individuals being born with the disease yearly [2]. The most predominant and severe type of SCD is caused by the homozygous inheritance of HbS (HbSS disease, or sickle cell anemia) [3]. SCD may also originate from co-inheritance of HbS with other hemoglobin mutations, such as hemoglobin C (HbSC) or β-thalassemia (HbSβ-thalassemia) [4]. β-thalassemia leads to absent (β^0^) or reduced (β^+^) synthesis of the beta-globin chains, and the coinheritance of HbS with β-thalassemia can be further categorized into two forms: HbSβ^+^ thalassemia and HbSβ^0^ thalassemia [5]. The genotypes HbSC and HbSβ^+^ tend to result in milder phenotypes, while HbSβ^0^ thalassemia may be as severe as HbSS disease [6].

Neurological complications of SCD include increased risk of stroke, silent cerebral infarction, and cognitive impairment, and are more common in individuals with HbSS disease [7, 8]; pediatric stroke, in particular, happens almost exclusively in individuals with HbSS. HbSS individuals also tend to display lower cognitive performance when compared with individuals with HbSC [9], possibly because anemia is less pronounced in HbSC. However, little research has been done on the impact of HbSC on the brain in adults.

Reduced cortical and subcortical volume has been previously reported in patients with SCD when compared with healthy controls. Kirk et al [10] showed cortical thinning in multiple brain regions in patients with SCD ranging from 5 to 21 years-old. They hypothesized that cortical thinning could be related with impaired cerebral perfusion. Mackin et al. [8] reported reduced cortical thickness in the temporal lobe in adults with SCD, while Kawadler et al. [11] found significantly reduced volume in the hippocampus of SCD pediatric patients with silent cerebral infarction. Overall, evidence of the impact of SCD on brain regions remains limited, particularly in adults.

The hippocampal involvement in higher cognitive processing has been well investigated, especially as pertains the hippocampal primary role in short-term and episodic long-term memory and learning [12]. The hippocampus is a bilaminar brain structure situated in the medial temporal lobe and it is comprised of the Cornu Ammonis (CA) – which is further divided into three conventional histological subregions, CA1-CA3 – and the Dentate Gyrus (DG) [13]. Additionally, the hippocampal formation includes the Subiculum (Sub) and the Entorhinal Cortex (ErC) [14, 15]. Hippocampal subfields are functionally interconnected, however functionalities are known to vary among the different subfields. For example, DG is suggested to be a preprocessor of incoming information, preparing it for subsequent processing in CA3, and potentially mediating learning, memory, and spatial encoding [16]. ErC, on the other hand, serves as the major input-output structure and mediates hippocampal connectivity with cortical regions [17]. Cells in the DG receive excitatory input from ErC and send excitatory output to the CA3 region via the mossy fibers [16]. Also, CA1 and CA3 are believed to contribute to episodic memory processing [18, 19]. CA2 is the smallest subfield but studies found that lesions in this area can lead to abnormal social behavior by impairing social recognition memory [20].

As pathology within the hippocampus is increasingly linked to a number of neurological and neuropsychiatric diseases such as Alzheimer’s disease, epilepsy, and depression [21-25], interest in the subfields of the hippocampus has grown. Subregions of the hippocampus are differentially affected, depending on the disease [26]. Recent advances in MRI techniques, such as 7 Tesla (T) MRI, have improved the spatial resolution and signal-to-noise ratio of MRI images [27]. These improvements enable more accurate volumetric analysis of the hippocampus subfields in *in vivo* studies, and improve automatic segmentation methods [28-32]. Manual segmentation of hippocampal subregions is time-consuming and labor intensive, limiting data collection and processing in large studies [23, 31]. To compensate for this shortcoming, automated segmentation methods have been developed [29, 31, 33].

This study aims to investigate how SCD impacts the volumes of hippocampus subfields. We used 7T MRI and an innovative radiofrequency (RF) coil to acquire high-resolution images of the hippocampal formation from a cohort of SCD participants and controls (N=77). 7T tailored preprocessing and automatic segmentation software [29] were used to compute the volumes of the hippocampus subfields. We hypothesized that the participants with SCD would exhibit lower hippocampal volume when compared with the control group and that the disease could affect specific hippocampal subregions.

## Methods

### Participants

Patients with SCD were recruited from the University of Pittsburgh Medical Center (UPMC) Adult Sickle Cell Program outpatient clinic under the University of Pittsburgh IRB protocols PRO12040139 and PRO08110422. Data were collected from 53 patients and 47 controls as part of a longitudinal study of neuroradiological biomarkers in SCD (ClinicalTrials.gov Identifier: NCT02946905). All patients with HbSS, HbSC and HbSβ thalassemia older than 18 years old and able to provide informed consent were informed about the study by staff members during their routine clinic visit, and offered entry into the study if they were in steady-state SCD (i.e. two weeks from an acute illness, including vaso-occlusive pain episodes). Eligibility criteria also included: 1) English-speaking; and 2) currently receiving routine follow-up care at the UPMC Adult Sickle Cell Program. Exclusion criteria included: 1) pregnancy, as determined by a positive urine human chorionic gonadotropin test at the time of the of the MRI; 2) acute medical problems including, but not limited to, acute vaso-occlusive episodes. Healthy controls, included in ClinicalTrials.gov identifier NCT02946905, were recruited from the community via brochures and from the University of Pittsburgh Clinical and Translational Science Institute Pitt+Me Registry if older than 18 and able to provide informed consent. Healthy controls were age, race, and gender matched to the participants with SCD. Both patients with SCD and controls were excluded if pregnant or lactating, had any medical condition that could result in neurocognitive or brain dysfunction (other than those resulting from SCD), including diabetes mellitus, coronary artery disease, peripheral vascular disease, and other causes of cerebral vasculitis such as Systemic Lupus Erythematosus (SLE), or contraindications to MRI scanning such as electronic implants, magnetically-activated implants, tattoos above the shoulders, or brain implants.

### MRI acquisition

MRI images were acquired using a 7T human MRI scanner (Siemens Magnetom, Germany) with a customized radiofrequency (RF) head coil with 16 transmit channels and 32 receive channels [34]. This RF coil produces homogenous images and whole brain coverage at 7T [27, 35-38]. T1-weighted (T1w) Magnetization Prepared RApid Gradient Echo (MPRAGE) was acquired with 0.75 mm isotropic resolution and the following parameters: TE/TI/TR = 2.17/1200/3000 ms, bandwidth = 391 Hz/Px, acceleration factor (Grappa) = 2, time of acquisition = 5:02 min. T2w images were acquired with high resolution in the plane perpendicular to the main axis of the hippocampus, slightly slanted from coronal acquisition. The sequence utilized was 2D turbo spin echo (TSE) with the following parameters: resolution = 0.375 x 0.375 x 1.5 mm^3^, TE/TR = 61/10060 ms, bandwidth = 264, acceleration factor (Grappa) = 2, time of acquisition = 3:32 min. Gradient echo (GRE) sequence – utilized as a secondary contrast for the calculations of the intracranial volume masks – was acquired with the following parameters: TE/TR = 8.16/24 ms, resolution 0.375 x 0.375 x 0.75 mm, acceleration factor 2, time of acquisition 8:20 min.

### Image preprocessing and hippocampus segmentation

Both T1w and T2w images were denoised using an optimal forward and inverse variance-stabilizing transformation (VST), specifically designed for the Rice distribution [39]. The forward VST converts the Rician heteroscedastic noise into a homoscedastic one (approximately Gaussian distributed), and thus the block-matching 4D (BM4D) denoising algorithm [40] was applied, followed by the inverse VST, to obtain the denoised images [41]. The images were then bias corrected with the SPM12 package using 30 mm of cutoff for the full width at half maximum (against 60mm of the default) [42]. **Figure 1** shows the preprocessing methods applied in a representative T2w image from this study. The bias correction and denoising methods improved homogeneity and mitigated noise in the images.

**Figure 1:**
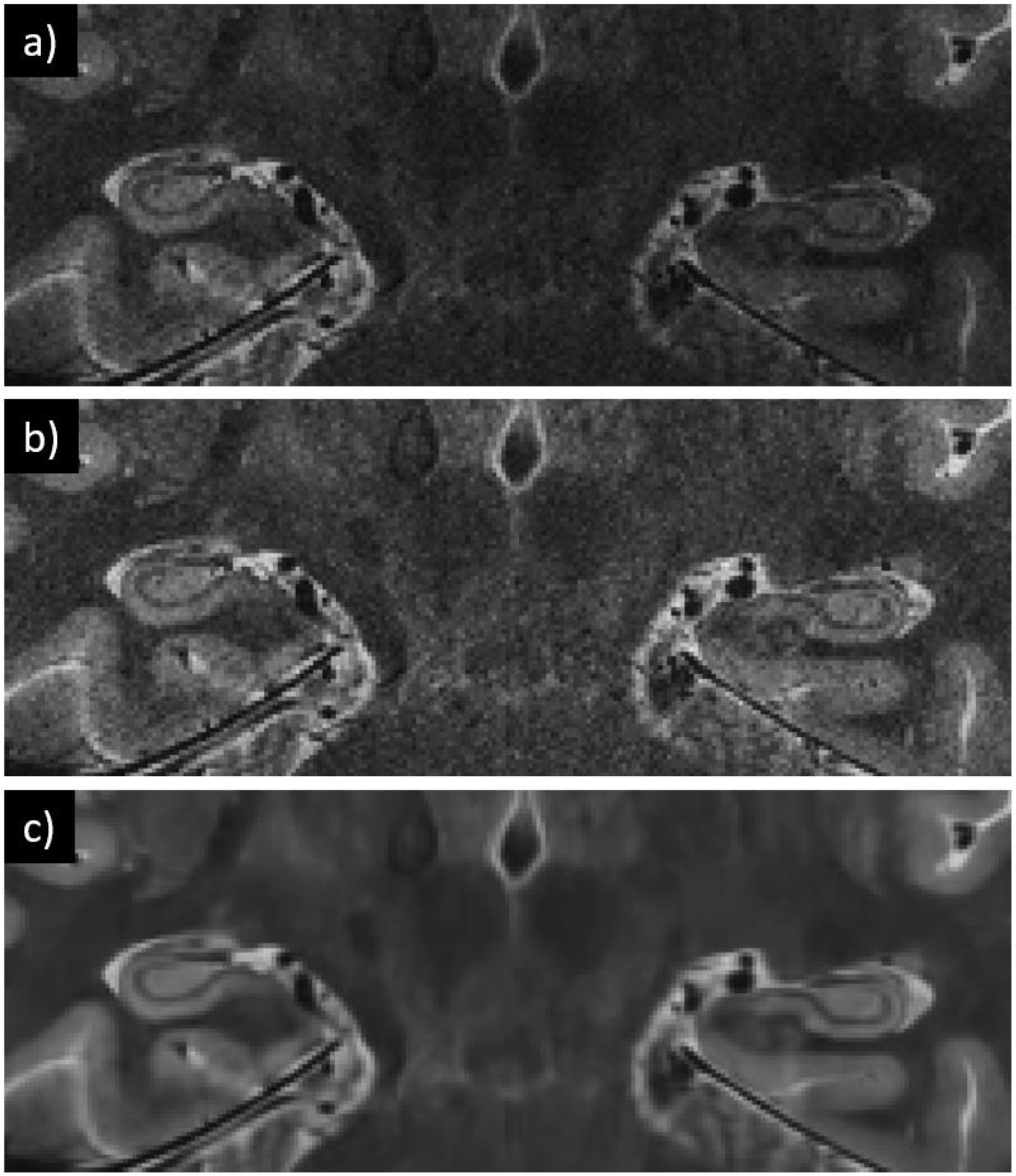
Preprocessing of the T2-weighted images in a representative SCD patient. In a), raw image from the 7T scanner. In b), bias corrected image. In c), bias corrected and denoised image

The automatic segmentation of hippocampal subfields (ASHS) package [29] and a 7T atlas from young adults’ data [43] were utilized to segment hippocampal subfields. The images from the atlas were denoised using the same technique described above for the T1w and T2w images. The results of the segmentation were manually corrected when the fixes were obvious and following the strategies described by Berron et al. [43]. The corrections were performed using the software ITK-SNAP [44]. In 23 of the 100 subjects, either the automatic segmentation failed or the manual fixes were not obvious (mostly due to excessive motion artifact); these subjects were excluded from the analysis, yielding a final sample size of 77. **Figure 2** shows the hippocampus segmentation in a representative SCD patient. The preprocessing techniques applied before segmentation with the ASHS package produced a smoother and more accurate segmentation of the hippocampus subfields. A summary of the methods is shown in **Figure 3**.

**Figure 2:**
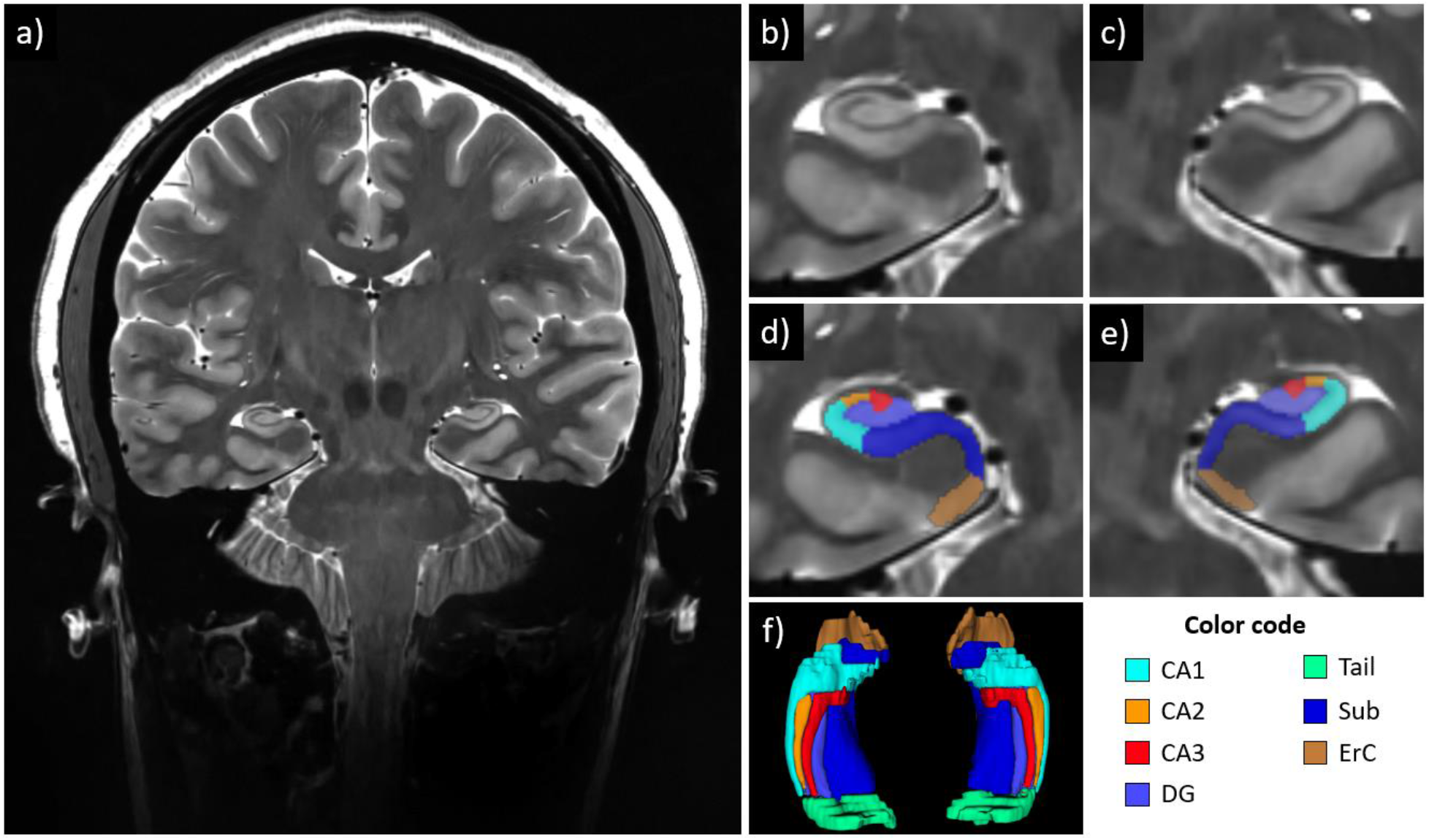
Example of hippocampal subfields segmentation in a patient with SCD. a) coronal slice of the T2-weighted image; b,c) zoomed image showing details of the hippocampus structure, subject right and left, respectively; d,e) hippocampus subfields segmentations overlaying the T2-weighted image, subject right and left, respectively; f) 3D reconstruction of the hippocampal subfield segmentations. Abbreviations - cornu ammonis 1-3, CA1-3; dentate gyrus, DG; hippocampal tail, Tail; subiculum, Sub; entorhinal cortex, ErC.

**Figure 3:**
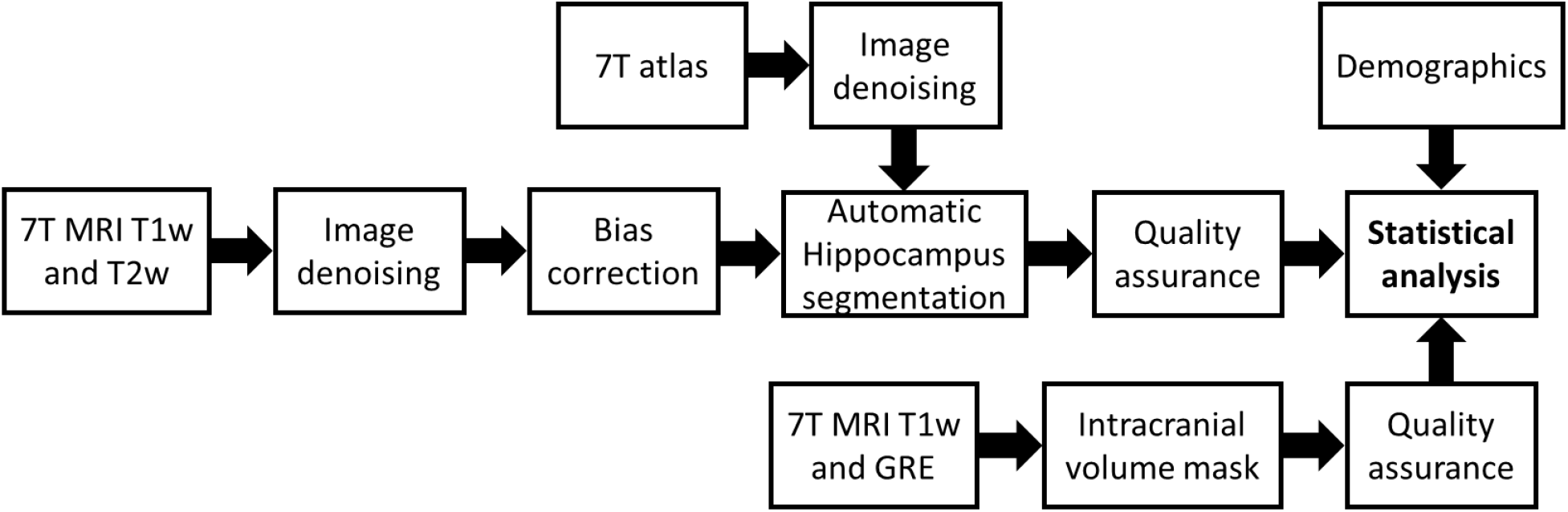
Flowchart of the method implemented for the hippocampal subfields segmentation and analysis. T1w, T1-weighted contrast; T2w, T2-weighted contrast; GRE: gradient echo sequence.

Intracranial volume is often used as a covariate in brain volumetric analysis; intracranial masks were calculated with the SPM12 package using the MPRAGE and GRE acquisitions as channels for the segmentations after a co-registration. The segmentations for the white matter, gray matter, and cerebrospinal fluid were combined, and the missing pixels in the segmentation were filled using the MATLAB function *imfill*. The imperfections in the segmentations were manually corrected using the ITK-snap software [44].

### Statistical Analysis

Sample characteristics were compared between SCD groups and controls using one-way ANOVA. The volumes of the hippocampal subfields were compared for patients and controls using analysis of covariance (ANCOVA), with gender, age, and intracranial volume as covariates. Those measures that differed by SCD status (patients and controls) were considered for inclusion in multivariable models testing the associations of SCD status and neuroimaging measures. The p-value threshold for between-group differences was calculated based on the Bonferroni correction. The statistical analysis was performed using SPSS (version 26, IBM, Chicago, IL).

## Results

### Demographics and hemoglobin values of the participants

There was no statistically significant difference in the age or gender between the groups (**Table 1**). As expected, the hemoglobin levels were significantly different between groups, with the severe SCD group (HbSS and HbSβ^0^) having the lowest Hb level (9.10 ± 1.09 g/dL), the milder SCD group (HbSC and HbSβ^+^) having an intermediate level (11.25 ± 2.48 g/dL), and the controls having the highest Hb level (13.94 ± 1.55 g/dL) at the time of the assessment (**Table 1**).

**Table 1.** Sample Characteristics

|  | ***SCD HbSS, HbS$\beta^0$***<br>***(severe)*** | ***SCD***<br>***HbSC, HbS$\beta^+$***<br>***(milder)*** | ***Controls*** | ***F-statistic (p-***<br>***value)*** |
| --- | --- | --- | --- | --- |
| *N* | 16 | 21 | 40 | - |
| *Age, mean (SD)* | 34.31(10.23) | 35.29(12.24) | 36.15(13.24) | 0.131 (0.88) |
| *Gender, F/M* | 7/9 | 9/12 | 16/24 | 0.41 (0.96) |
| ***Hb<sup>1</sup>, mean (SD)*** | **9.10 (1.09)** | **11.25 (2.48)** | **13.94 (1.55)** | **43.5 (6.9E-13) \*** |
| <sup>1</sup> in g/dL |  |  |  |  |
| * **p-value < 0.05** |  |  |  |  |

### Neuroimaging characteristics of patients with SCD vs. controls

Results of the comparison of the hippocampal subfields volumes between SCD patients and controls are tabulated in **Table 2. Figure 4** shows the bar plots with the individual volumetric data points for all the subjects included in this study. Significant differences between the hippocampal volumes of the SCD and control groups were observed bilaterally in the region that encompasses the DG, CA2, and CA3: - 11.55% (F = 20.79, p = 0.020×10^−3^) in the left hippocampus, and - 11.36% (F = 14.09, p = 0.350×10^−3^) in the right hippocampus. There was also a trend towards a reduction of the left CA1 (−10.52%, F = 5.69, p-value 0.020), right CA1 (−8.02%, F = 4.02, p-value 0.049), and left ErC (−9.03%, F = 5.68, p-value 0.020), which was not statistically significant after Bonferroni correction. The other hippocampal subregions, including the right ErC, bilateral Tail, and bilateral Sub, did not significantly differ between SCD and control groups. The whole left (−7.29%, F = 7.29, p-value 0.009) and right hippocampus (−6.57%, F = 4.93, p-value 0.030) showed a trend that did not remain statistically significant after Bonferroni correction. Between-group differences in the region containing DG, CA2, and CA3 were marginally attenuated after adjustment for intracranial volume (from 11.55% to 10.10% for the left hemisphere, and from 11.36% to 9.63% for the right hemisphere), and minimally attenuated with the inclusion of age and gender in the model, as shown in **Table 3**. No significant differences were found between the severe and milder SCD genotypes.

**Table 2.** Neuroimaging Characteristics. Mean volume in mm^3^, SD and percent difference

|  | **SCD**<br>**(severe and**<br>**milder)**<br>**--**<br>**Mean (SD)** | **Controls**<br>**--**<br>**Mean (SD)** | **Percentage**<br>**difference**<br>**SCD vs**<br>**controls<sup>5</sup>** | **F-statistic (p-value)** |
| --- | --- | --- | --- | --- |
| *Intracranial volume<sup>1</sup>* | 1306040<br>(114740) | 1336886<br>(130234) | -2.31% | 2.813 (0.097786)<sup>a</sup> |
| *Total hippocampal volume<sup>1</sup>* | 4635.69 (541.56) | 4980.17<br>(557.54) | -6.92% | 6.400 (0.013601)<sup>b</sup> |
| *Asymmetry Ratio<sup>2</sup>* | 0.03655 (.02280) | 0.03385<br>(.02242) | 7.97% | 0.475 (0.492749)<sup>b</sup> |
| **Left Hemisphere** |  |  |  |  |
| *Cornu Ammonis 1<sup>1</sup>* | 619.02 (115.08) | 691.82 (122.65) | -10.52% | 5.69 (0.019689)<sup>b</sup> |
| ***Dentate Gyrus<sup>1,3</sup>*** | **632.49 (70.68)** | **715.05 (92.82)** | **-11.55%** | **20.79 (0.000020)<sup>b\*</sup>** |
| *Subiculum<sup>1</sup>* | 979.63 (116.29) | 999.65 (133.50) | -2.00% | 0.02 (0.899994)<sup>b</sup> |
| *Entorhinal cortex<sup>1</sup>* | 743.54 (133.47) | 817.36 (131.31) | -9.03% | 5.68 (0.019773)<sup>b</sup> |
| *Tail<sup>3</sup>* | 583.22 (123.34) | 561.99 (96.57) | 3.78% | 1.34 (0.251466)<sup>b</sup> |
| *Hippocampus<sup>1</sup>* | 2231.14 (250.92) | 2406.52<br>(281.49) | -7.29% | 7.29 (0.008650)<sup>b</sup> |
| **Right Hemisphere** |  |  |  |  |
| *Cornu Ammonis 1<sup>1</sup>* | 685.96 (116.09) | 745.73 (108.93) | -8.02% | 4.02 (0.048854)<sup>b</sup> |
| ***Dentate Gyrus<sup>3</sup>*** | **669.81 (94.72)** | **755.69**<br>**(103.89)** | **-11.36%** | **14.09 (0.000350)<sup>b\*</sup>** |
| *Subiculum<sup>1</sup>* | 1048.77 (145.29) | 1072.22<br>(131.94) | -2.19% | 0.01 (0.918181)<sup>b</sup> |
| *Entorhinal cortex<sup>1</sup>* | 799.98 (186.52) | 841.61 (124.96) | -4.95% | 0.81 (0.371777)<sup>b</sup> |
| *Tail<sup>1,4</sup>* | 499.13 (84.39) | 494.83 (69.95) | 0.87% | 0.46 (0.501763)<sup>b</sup> |
| *Hippocampus<sup>1</sup>* | 2404.55 (300.00) | 2573.64<br>(287.13) | -6.57% | 4.93 (0.029504)<sup>b</sup> |
| <sup>1</sup> in mm<sup>3</sup><br><sup>2</sup> Defined as (right hippocampus - left hippocampus)/(right hippocampus + left hippocampus)*100<br><sup>3</sup> Includes Dentate Gyrus, Cornu Ammonis 2 and 3<br><sup>4</sup> Defined in the reference [23]<br><sup>5</sup> Given 95% confidence interval<br><sup>a</sup> Corrected for age and gender<br><sup>b</sup> Corrected for age, gender, and intracranial volume<br>**\* Significantly different after Bonferroni correction (p-value threshold: 0.05 / 10 subfields / 2 groups = 0.0025 for significance)** |  |  |  |  |

**Table 3.**
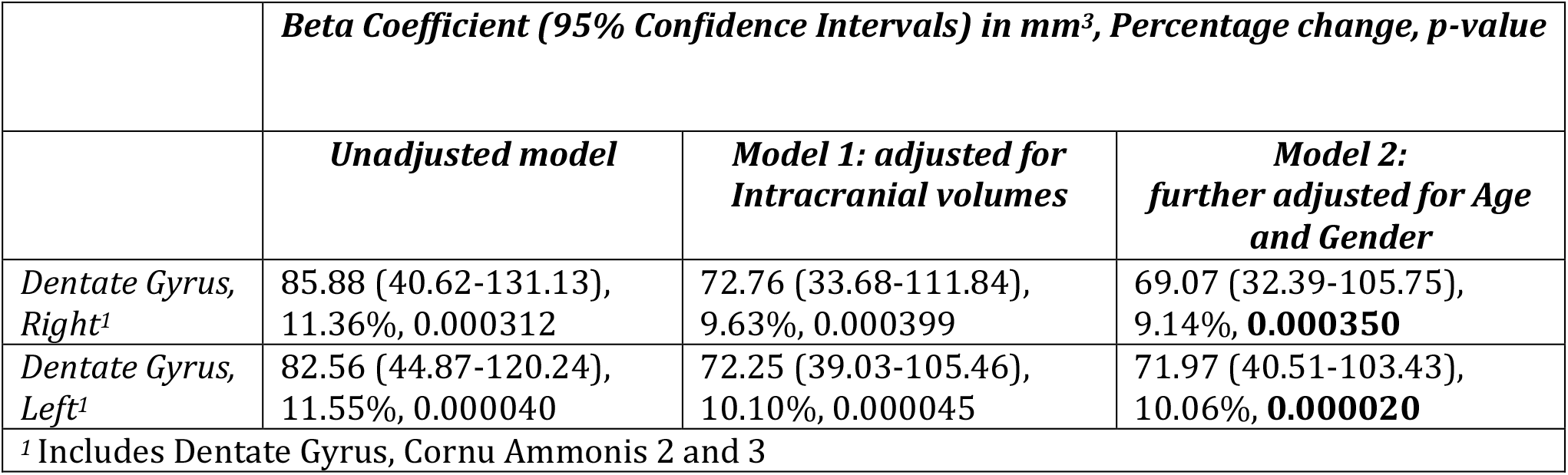
Multivariable Regression Models of Sickle Cell Disease Status predicting Dentate Gyrus volume

**Figure 4:**
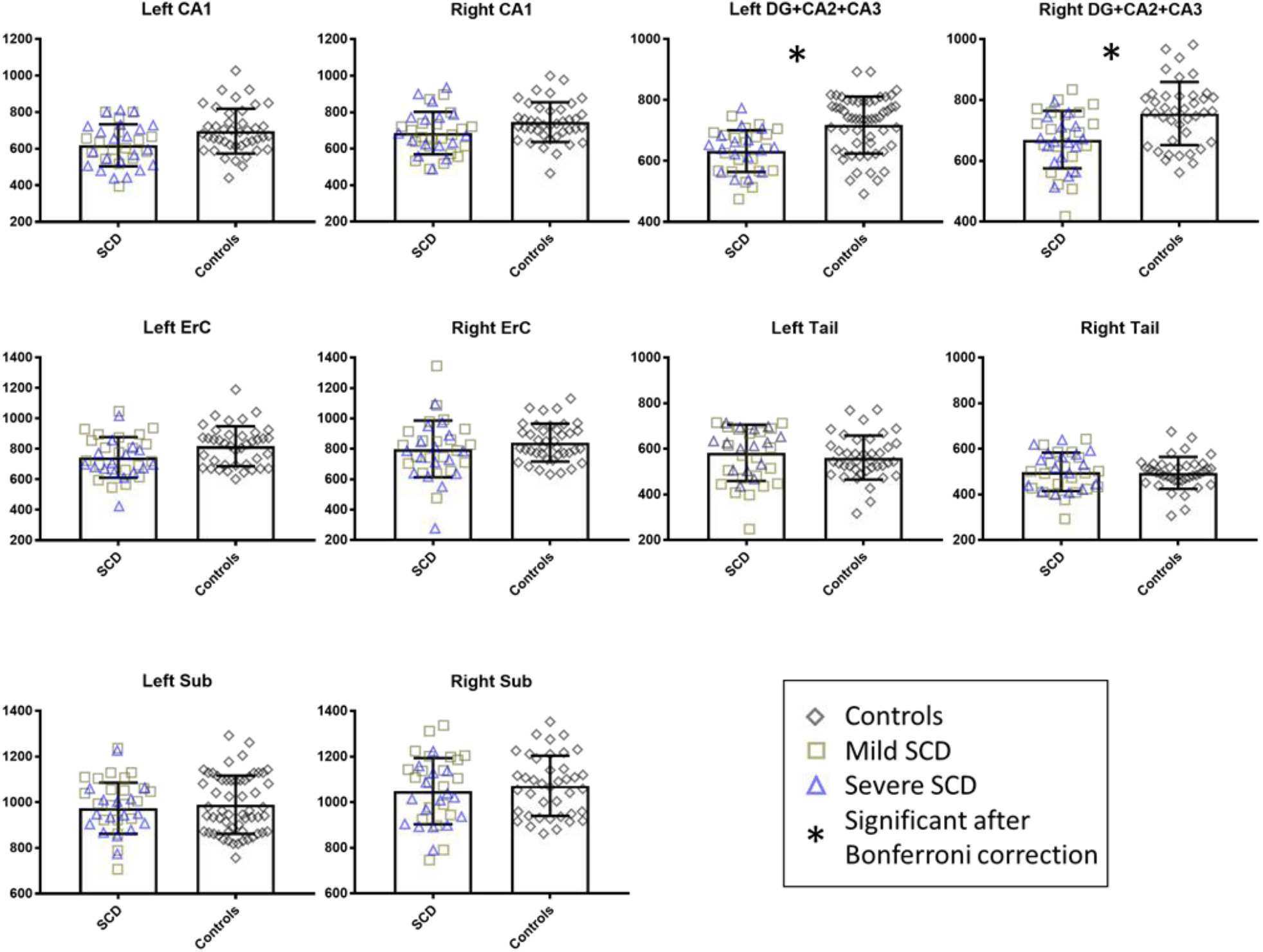
Hippocampal subfield volume comparison between patients and controls. There was a significant volume difference between the patient and the control groups in the region formed by the Dentate Gyrus (DG), CA2, and CA3 bilaterally. Abbreviations - cornu ammonis 1-3, CA1-3; dentate gyrus, DG; entorhinal cortex, ErC; hippocampal Tail, Tail; subiculum, Sub. The error bars represent the standard deviation of the data. Controls (n=40), Milder SCD (n=21), Severe SCD (n=16).

## Discussion

In this study, 7T MRI and innovative radiofrequency coil developments allowed the acquisition of high-resolution and high-quality images of the hippocampus. Preprocessing techniques such as denoising and bias correction further improved the image quality. Manual quality assurance and correction refined the automatic hippocampal segmentation method. The use of the GRE sequence with the T1-weighted MPRAGE also improved the estimation of the intracranial volume masks, although manual corrections were still necessary.

Compared with controls, our analysis shows that the region DG+CA2+CA3 is the most affected hippocampal subregion in the SCD group, an effect observed with a mean difference of −11.55% on the left hemisphere and −11.36% on the right hemisphere. After adjusting for age, gender, and intracranial volume, the differences were marginally reduced, indicating that SCD is the major contributor to the overall difference. Moreover, the hippocampal subregions CA1 bilaterally, left ErC, and hippocampus bilaterally also showed a trend towards smaller volumes in the SCD group, but the differences were not statistically significant after Bonferroni correction.

These results are consistent with findings in pediatric populations with SCD. Kawadler et al. [11] reported a volume difference of −6.74% in the right hippocampus and −10.26% in the left hippocampus in pediatric HbSS subjects with silent cerebral infarction when compared with healthy controls. If confirmed in future studies, our results indicate that the impact of SCD on hippocampal morphology reported in pediatric patients persists in adulthood. Future studies should assess the impact of these patterns of significant subregional hippocampal atrophy on cognitive decline.

To the best of our knowledge, our study is the first to analyze hippocampal subregions with high-resolution images obtained with 7T MRI, and to include both severe and milder genotypes of SCD. Understanding how hippocampal subregion volumes differ in SCD may identify biomarkers of disease severity and cognition. For instance, our analysis found that the total hippocampal volume was not significantly different between patients and controls, whereas the DG+CA2+CA3 subregion grouping showed a consistent volumetric difference between the two groups. This hippocampal region is believed to mediate learning, memory, and spatial encoding [16], and abnormalities in this area could be associated with deficits in these cognitive domains in SCD.

The hippocampus is particularly vulnerable to hypoxia and inflammation [45], which are common pathogenic mechanisms in SCD. Mouse models of SCD reveal that hippocampal pathology is associated with cognitive deficits [46]. Interestingly, we did not observe an association between the volume of hippocampal subregions and SCD genotype, despite evidence that individuals with milder genotypes tend to have better cognitive functioning [47, 48]. It is possible that compensatory mechanisms such as increased hippocampal activation and connectivity [49, 50] may confound the effect of hippocampal atrophy on cognition.

One limitation of this study is that hippocampal subregions may be inaccurately labeled as the adjacent subregions. In particular, CA2 and CA3 subregions are small and their borders do not present clear landmarks in MRI; therefore, they are more prone to mislabeling errors. On the other hand, the segmentation of DG is the most accurate [29], since it follows well defined landmarks in the T2-weighted image. Nevertheless, only histological data could precisely identify all specific subregions. To reduce the impact of mislabeling, we combined CA2 and CA3 with DG, a common strategy [51]. To further enhance the precision of our measurements, we excluded 23 subjects whose scans presented excessive motion artifacts, mostly due to artifacts in the T2w TSE acquisition. Future work in RF coil design, MRI sequence development, and image processing may lead to a higher inclusion rate.

In summary, we found that between-group differences in the hippocampus followed a distinct spatial distribution, being more pronounced in the DG+CA2+CA3 bilaterally as compared to other subregions. These associations were attenuated after adjustment for intracranial volume and demographics, but remained significant. Further studies will be necessary to clarify the mechanisms that lead to volume reduction in the hippocampal subfields and elucidate their significance as an imaging biomarker for cognitive deficits in individuals with SCD.

## Acknowledgments

This work was supported by the National Institutes of Health under award numbers: R01HL127107, R01MH111265, R01AG063525, and T32MH119168. The first author was partially supported by CAPES Foundation, Ministry of Education of Brazil, under the award number 13385/13-5. This research was also supported in part by the University of Pittsburgh Center for Research Computing (CRC) through the resources provided.

